# The Israeli Acute Paralysis Virus IRES captures host ribosomes by mimicking a ribosomal state with hybrid tRNAs

**DOI:** 10.1101/606236

**Authors:** Francisco Acosta-Reyes, Ritam Neupane, Joachim Frank, Israel S. Fernández

## Abstract

The <u>C</u>olony <u>C</u>ollapse <u>D</u>isorder or CCD is a multi-faceted syndrome decimating bee populations worldwide[1]. A group of viruses of the widely distributed *Dicistroviridae* family have been identified as a causing agent of CCD[2]. This family of viruses employ non-coding RNA sequences, called <u>I</u>nternal <u>R</u>ibosomal <u>E</u>ntry <u>S</u>ite (IRES), to precisely exploit the host machinery for protein production. Using single-particle cryo-electron microscopy (cryo-EM) we have characterized at high resolution how the IRES of the intergenic region of the <u>I</u>sraeli <u>A</u>cute <u>P</u>aralysis <u>V</u>irus (IAPV) captures and redirects translating ribosomes towards viral messengers. Through a series of six structures at nominal resolutions close to 3Å, we could reconstruct the trajectory of IAPV-IRES from an early small subunit recruitment to a final post-translocated state in the ribosome. An early commitment of IRES/ribosome complexes for global pre-translocation mimicry explains the high efficiency observed for this IRES. The presented structures will help guide on-going efforts directed towards fighting CCD through RNA-interference technology [3].

## Introduction

*Apis mellifera*, the common western bee, is affected worldwide by an enigmatic syndrome characterized by a drastic disappearance of the workforce, causing the accelerated collapse of the hive[1]. Given the essential role bees play in pollination of economically important crops, the impact of this syndrome, termed <u>C</u>olony <u>C</u>ollapse <u>D</u>isorder or CCD, has been estimated to cost the US economy $15 billion in direct loss of crops and $75 billion in indirect losses [4].

Though the exact etiology of CCD is unknown [5], a group of viruses belonging to the *Dicistroviridae* family were found in metagenomic studies of CCD affected hives [2, 6]. Among this group of viruses, the <u>I</u>sraeli <u>A</u>ccute <u>P</u>araysis <u>V</u>irus (IAPV) showed a strong correlation with CCD, revealing a prominent role in the developing of the syndrome [7, 8].

The *Dicistroviridae* family of viruses exhibits a wide environmental distribution, targeting invertebrates, mainly insects and arthropods [9]. The genetic architecture of these viruses is composed of a single RNA molecule in the positive sense ((+)-RNA)) which contains two open reading frames (ORF1 and ORF2) [10, 11]. ORF1 encodes nonstructural proteins: a RNA helicase, a cysteine protease and a RNA-dependent RNA polymerase (RdRP). The second ORF encodes a single poly-protein that, upon proteolytic digestion, generates the structural proteins that will eventually compose the viral capsid [12].

Both ORFs are flanked by non-coding RNA sequences responsible for the regulation of the expression of their downstream genes [10, 13, 14]. A fine balance between the expression of ORF1 and ORF2 is required for the replication and expansion of the virus [15, 16]. This is achieved through a precise exploitation of host resources, specially the machinery for protein synthesis [12]. The flanking non-coding RNA regions preceding both ORFs harbor two different <u>I</u>nternal <u>R</u>ibosomal <u>E</u>ntry <u>S</u>ites (IRES) [10]. IRESs are structured RNA sequences able to interfere with canonical translation, capturing host ribosomes in order to redirect them towards the production of viral proteins [17].

Eukaryotic ribosomes are operated by a complex collection of cellular factors that regulate the production of proteins in the cell [18]. Specially regulated in eukaryotes is the first step of translation, initiation [19]. Along this initiation phase, the small ribosomal subunit (40S), in partnership with many initiation factors, is able to capture a mRNA, localize its AUG initiation codon, deliver the first aminoacyl-tRNA and finally recruit the large subunit (60S) in oder to assemble an elongation competent ribosome (80S) primed with an aminoacyl-tRNA in the P site and a vacant A site [20].

The majority of IRES families leverage the complexity of initiation to hijack cellular ribosomes [17, 21]. However, the IAPV-IRES found in the intergenic region of the IAPV virus belong to the well characterized type IV family of viral IRESs. This group of IRES sequences dispense with all canonical initiation factors and is able to assemble by themselves an elongation competent ribosome, successfully redirecting the cellular machinery for protein production by a RNA-only mechanism[22]. This is accomplished by an elaborated use of intrinsically dynamic elements of the ribosome, naturally involved in translocation [23]. These IRESs are able to induce an artificial state on the ribosome, mimicking a pre-translocation state with tRNAs. Elongation factors eEF2 and eEF1A can then be recruited to effectively bypass the highly regulated initiation [24, 25, 26], jumpstarting directly in the elongation phase [27].

The type IV IRES family exhibits a remarkable structural diversity, which remains poorly characterized [22]. Two genera, based on phylogenetic analysis of ORF2 as well as in the intergenic region, can be defined: Aparaviruses and Cripaviruses. The <u>Cr</u>icket <u>P</u>aralysis <u>V</u>irus <u>IRES</u> (CrPV-IRES), a prototypical Cripavirus, has been extensively studied due to its early discovery and use as model mRNA of early studies in translation [28, 29]. Recently, a divergent IRES sequence of a shrimp infecting virus, the Taura Virus Syndrome IRES, has been visualized by cryo-EM in complex with yeast ribosomes [25, 30]. The IAPV-IRES presents the prototypical features of an Aparavirus, with an additional stem loop (SL-III) nested within the pseudo-knott I (PKI) and an extended L1.1 region [31]. Importantly, this IRES can drive translation in two different ORFs, able to produce two different polypeptides from the same mRNA. A frameshift event at the first coding codon is responsible for this multi-coding capacity [32].

Research efforts directed towards finding the cause of CCD and developing strategies to prevent it are underway [3, 6]. RNA-interference has proved effective in protection against CCD. Directing double stranded RNAs complementary to the IRES of the intergenic region of the IAPV virus decrease the probability of hive collapse, preventing effectively the massive death of the workforce, guaranteeing the protection of the queen and thus the survival of the colony [33].

Using single particle cryo-electron microscopy (cryo-EM) we have characterized at high resolution how the IAPV-IRES redirect the host machinery for protein synthesis exploiting novel ribosomal sites. An early commitment of IRES/ribosome complexes towards global pre-translocation mimicry explains the high efficiency in ribosome highjacking observed for this IRES. These results may inspire structure-based rational designs for the fight against CCD by RNA-interference technology [6].

## Results

### Biochemical set-up and cryo-EM strategy

Previous biochemical and genetic studies of IAPV-IRES established the secondary structure scheme displayed in figure 1A [31]. The prototypical architecture of the type IV IRES family consisting of three nested pseudoknotts is extended by a 5’ terminal <u>S</u>tem <u>L</u>oop (SL-VI) proposed to play functional roles in the early positioning of the IAPV-IRES in the ribosome [34, 35]. Additionally, the genus Aparavirus is characterized by an extended PKI which contains an insertion of a large stem loop (SL-III, Fig. 1A, bottom). In the IAPV-IRES, SL-III consists on 8 Watson-Crick canonical base pairs and a terminal loop of 6 nucleotides. Notably, two unpaired adenine residues are placed in a strategic position at the core of the three-way helical junction connecting SL-III, the <u>A</u>nti codon <u>S</u>tem <u>L</u>oop (ASL)-like element of the PKI (residues 6546-6574) and the double helical segment connecting PKI and PKIII. A <u>V</u>ariable <u>L</u>oop <u>R</u>egion (VRL) bridges the mRNA-like element of PKI (residues 6613-6617) with the helical region connecting PKI and PKIII. This single-stranded RNA loop is poorly conserved in sequence, however, even though its role in IRES functioning remains enigmatic, biochemical experiments have proved its integrity is mandatory for productive IRES-driven translation [36].

**Figure 1:**
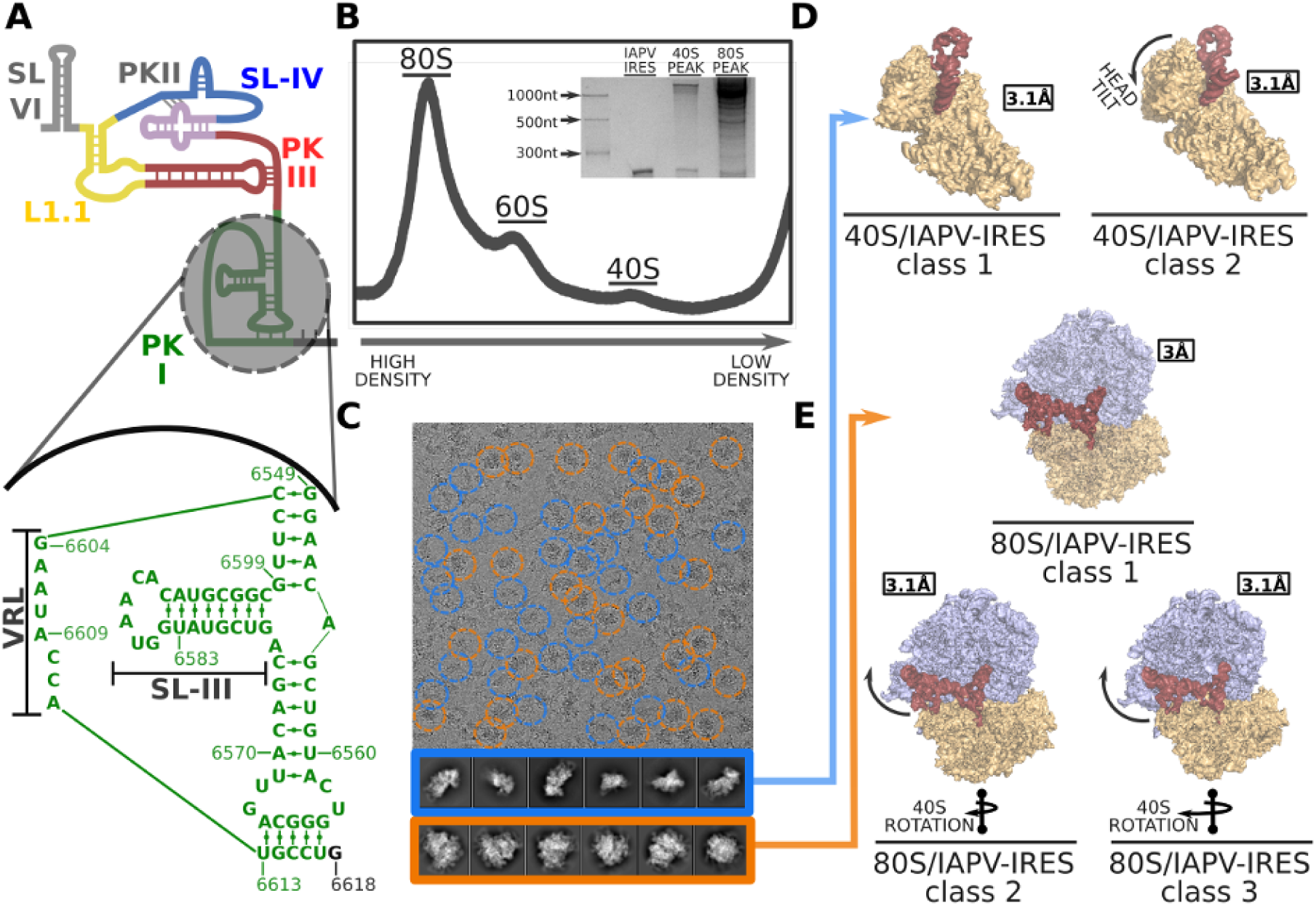
IAPV-IRES secondary structure, experimental set up and cryo-EM image processing workflow. (**A**) IAPV-IRES diagram colored according to secondary structure motifs. Bottom, a closer view of the IAPV-IRES PKI highlighting its sequence, with base pairs indicated as well as the Variable Loop (VRL) and stem loop III (SL-III). (**B**) Sucrose gradient UV profile of 60S/40S/IAPV-IRES reaction mixture after an overnight run. The peaks corresponding to 80S and 40S were used for RNA extraction and UREA-PAGE shown in the inset. (**C**) Representative cryoEM image where roughly half of the particles correspond to 40S (blue) and the other half to 80S (orange).(**D**) Two classes with robust density for the IAPV-IRES where found in the 40S group. (**E**) After classification, three classes with clear IAPV-IRES density and small differences in the conformation of the 40S where found in the 80S group.

In order to understand in structural terms how these constituents units of the IAPV-IRES are involved in ribosome hijacking, we produced a full, wild-type IAPV-IRES, including SL-VI and the first two coding codons. Binary complexes with mammalian ribosomes in complex with IAPV-IRES were generated incubating IRES with ribosomal subunits. We designed a reaction featuring double amount of 40S over 60S in an overall background of IRES excess, to test the ability of the IAPV-IRES to engage both 40S and full 80S ribosomes in a productive and stable binary interaction. The stability of these interactions was tested through a sucrose gradient run overnight (Fig. 1B) where the different complexes could be resolved according to their size differences. Each peak was subjected to RNA extraction and UREA-PAGE analysis where the presence of IAPV-IRES bound in the 80S peak as well as in the 40S peak could be confirmed (Fig. 1B).

Leveraging latest cryo-EM maximum likelihood classification methods implemented in RELION 3.0 [37, 38], we decided to directly image the preparation assayed for sucrose gradient analysis in cryo-EM grids, without further purification. A large dataset ensuing from this experiment was subjected to an optimized classification scheme combining different *in silico* classification approaches (Fig. S1) what allowed us to identify and refine to high resolution five distinctive classes from a single dataset (Fig. 1C-E).

### The IAPV-IRES restricts the conformational freedom of the 40S blocking functional sites

The small ribosomal subunit can be roughly divided in two parts: the body, which forms the bulk of the subunit accounting for two thirds of its mass, and a mobile part roughly comprising the remaining third, designated as the head (Fig. 2A). The interface between these two components forms the tRNA binding sites of the small subunit. The head of the 40S subunit is a dynamic component, modifying its relative orientation respect the body. This dynamic is of critical importance in two aspects of translation: the positioning of the initiator aminoacyl-tRNA and in the concerted movement of mRNA and tRNAs along elongation [39, 40]). We identified two classes of particles showing robust density for IAPV-IRES in the context of a binary interaction with the 40S (Fig. 2B and C). All elements of the IAPV-IRES included in the produced construct were identified in the maps except SL-VI, which proved disordered, no density could be assigned to it even in low-pass filtered maps. The L1.1 region in the context of a binary interaction with the 40S shows a high degree of mobility and can only be modeled in maps filtered to 4Å.

**Figure 2:**
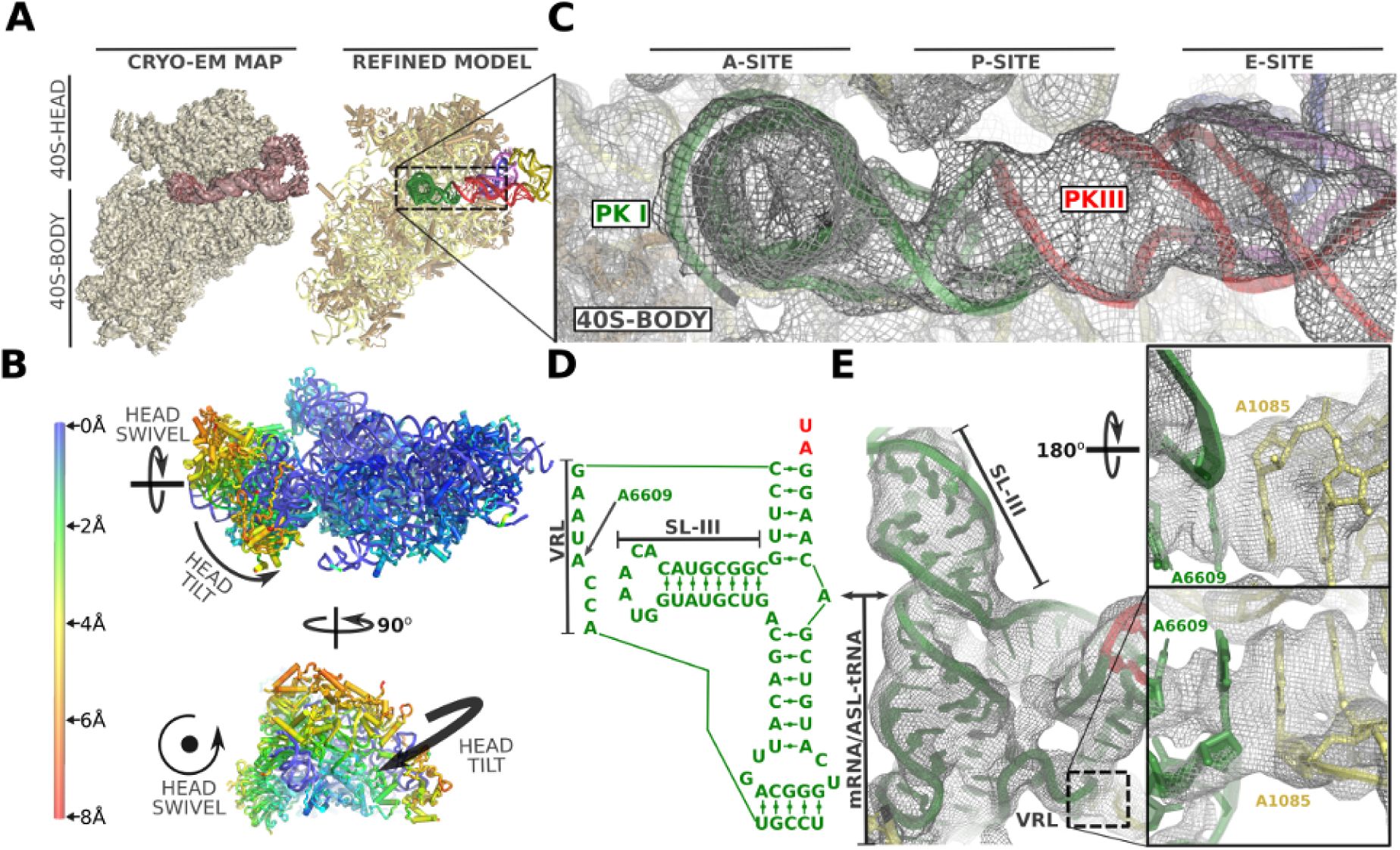
Structure of the IAPV-IRES in complex with the 40S ribosomal subunit. (**A**) Overview of the mammalian 40S in complex with IAPV-IRES. Left, cryo-EM final post-processed map of class 1 with 40S colored yellow and IAPV-IRES maroon. Right, corresponding final refined model with IAPV-IRES colored according to figure 1A. (**B**) Ribbon diagram of the 40S colored by pairwise root-mean-square deviation displacements observed between the two IAPV-IRES/40S classes. The different position of the 40S head between both classes is a composition of swiveling and tilt movements (indicated by arrows in orthogonal views). (**C**) Close up view of the ribosomal sites of the 40S for IAPV-IRES/40S class 1 showing cryo-EM unsharpened final experimental cryo-EM density. (**D**) Sequence of the PKI three-way helical junction. (**E**) Cryo-EM density for this region of the IAPV-IRES in class 1. Right, close up view of the VRL region reaching the P site. A stacking interaction is established between IRES adenine 6609 and 18S rRNA residue A1085.

The IAPV-IRES inserts two elements of its structure between the head and the body of the 40S subunit, effectively restricting the dynamics of the 40S head to specific ranges of conformations. The ASL/mRNA mimicking part of the PKI is inserted in the decoding site (A site) of the small subunit stabilized by the decoding bases of the 18S rRNA A1824-A1825 and G626 (A1492, A1494 and G530 in *E.coli* [41]), inducing a decoding event [41]. SL-IV is deeply inserted in the interface of head and body, in the surroundings of the E site, clamped by stacking interactions stablished with residue A6498 of the IRES and tyrosine 72 from uS7 and arginine 135 from uS11 (Fig. 3D and E). In this conformation, the IAPV-IRES fully blocks all three tRNA binding sites of the 40S subunit, interfering with early steps of canonical initiation (Fig. 2C) [19].

**Figure 3:**
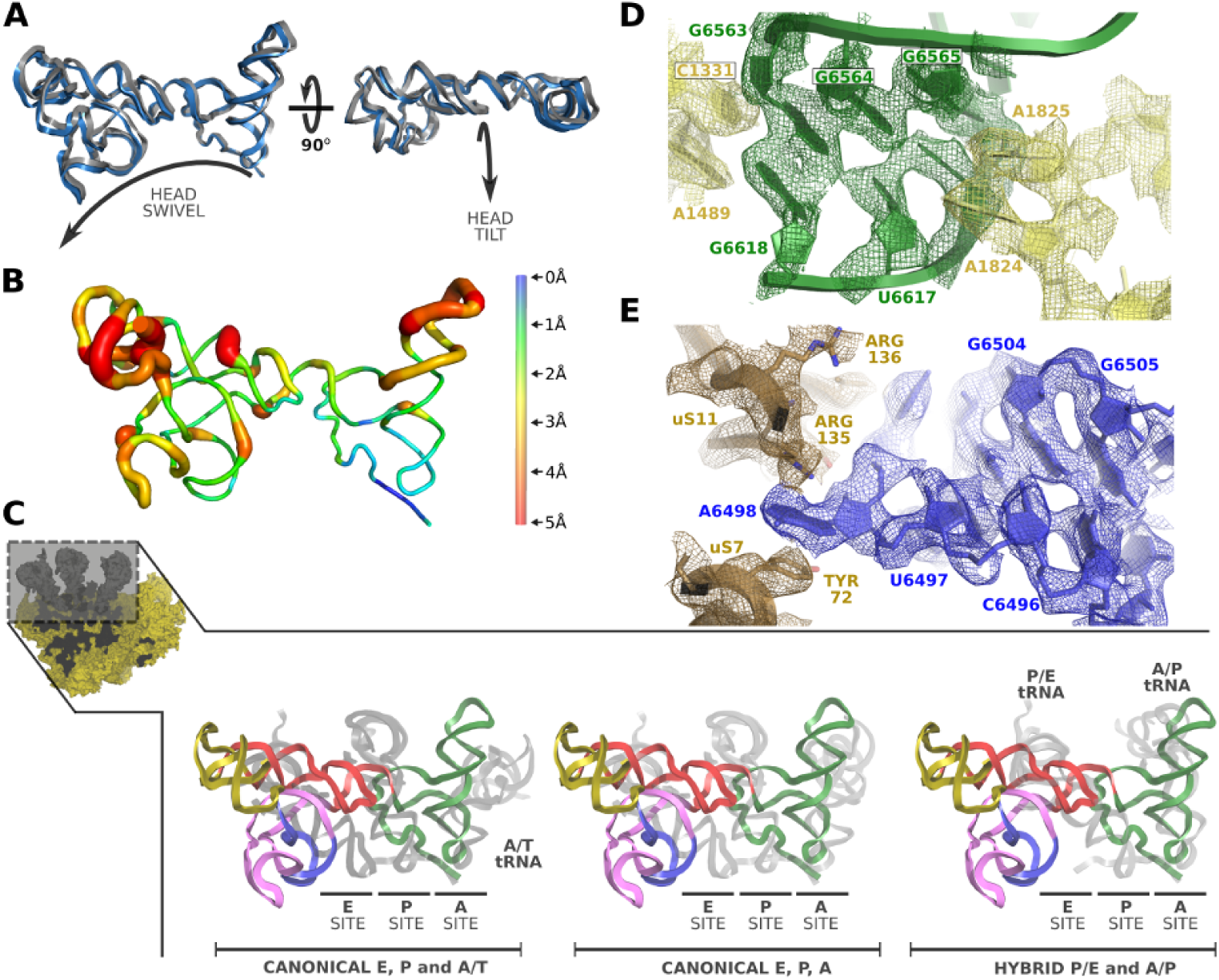
IAPV-IRES conformation in the context of a 40S interaction. (**A**) Superposition of IAPV-IRES models corresponding to IAPV-IRES/40S class 1 and 2 after alignments excluding the IRES and the 40S head. A similar conformation can be observed with distinctive relative orientation respects the 40S body. The movements characteristic of the 40S head are indicated by arrows in orthogonal views. (**B**) Ribbon diagram of the IAPV-IRES colored by pairwise root-mean-square deviation displacements observed between the two IAPV-IRES/40S classes. The ASL/mRNA like regions of the PKI and well as the SL-IV shows the lowest degree of displacement (blue) whereas the apical part of SL-III and the L1.1 the highest (red). (**C**) Superposition of the IAPV-IRES with tRNAs in different configurations indicated at the bottom. IAPV-IRES is depicted as ribbons colored according to the secondary structure elements and tRNAs are represented as grey ribbons. Alignments were computed with the 40S body, excluding from the computation the ligands (IRES/tRNAs) and the 40S head. (**D**) Detailed view of the refined model for IAPV-IRES/40S class 1 inserted in the final post-processed cryo-EM density focused on the decoding center of the 40S. PKI of IAPV-IRES is depicted green and 18S rRNA yellow. (**E**) Close up view of the refined model for IAPV-I9RES/40S class 1 inserted in the final post-processed cryo-EM density focused on the SL-IV of the IAPV-IRES (depicted blue).

Density for the full PKI, including SL-III, was clearly visible in the maps (Fig. 2E and S3) what allowed accurate modeling of the three-way helical junction characteristic of Aparavirus IRESs. The VRL was also visible in the maps, partially occupying the P site, stabilized by a stacking interaction between A6609 of the IRES and A1085 of the 18S rRNA (Fig. 2E,right).

The two classes present a conformation of the IAPV-IRES nearly identical (r.m.s.d.=1.12Å between the two IRES conformations) but in displaced position respects the 40S body (Fig. 3A and B). The IAPV-IRES seems to follow the movement of the 40S head, pivoting around the anchored PKI and SL-IV which are exceptionally stabilized by ribosomal elements from both head and body, effectively “clamping” the IRES to the 40S subunit (Fig. 3D and E).

The three-way helical junction modeled in the PKI of the IAPV-IRES resembles a “hammer” shape, with SL-III coaxially stacked on top of the ASL-like stem. Perpendicular to both and situated in between them, a helical segment connects PKI and PKIII. The coaxially aligned SL-III and ASL-like domain forms a straight unit of shape and dimensions similar to a tRNA, excluding the acceptor stem (Fig. 3C). Alignments of structures containing tRNAs in several configurations (canonical tRNAs PDBID:4V5D[42], with A/T-tRNA PDBID:5LZS[43] and hybrid tRNAs PDBID:3J7R[44]) with the structure of the IAPV-IRES in complex with the 40S, reveal an interesting positioning of the straight unit formed by SL-III and the ASL-like part of PKI (Fig. 3C). Notably, the SL-III/ASL-like unit of IAPV-IRES populates a space more similar to a hybrid A/P-tRNA than a canonical, A/A-tRNA or A/T-tRNA (following nomenclature of hybrid states previously proposed ([45]). Similarly, PKIII overlaps with the position occupied by a hybrid P/E-tRNA, mimicking its helical components the elbow region of a tRNA in this intermediate configuration. Overall, the IAPV-IRES is able to manipulate the 40S subunit in isolation, blocking the functional sites where key canonical initiation factors bind, and at the same time, steering the intrinsic dynamics of the 40S head towards a configuration reminiscent of an early elongation, pre-translocated state.

### SL-III interacts with ribosomal protein uL16 stabilizing the 80S in a pre-translocation configuration mimicking hybrid tRNAs

The binary IAPV-IRES/80S complex populates three mayor conformations, with limited differences between them (Fig. 1E). The majority of particles were assigned to a class where the 40S subunit exhibit a small degree of intersubunit rotation (aprox. 1°) compared to the unrotated, canonical configuration (Fig. 4A). No mayor swiveling or tilt of the 40S head is visible in this conformation. The IAPV-IRES maintains a similar global conformation as in the binary complex with 40S but, in the 80S map, both the L1.1 region as well as the tip of the SL-III shows good density due to their dynamics being restricted by specific contacts with elements of the 60S: the L1-stalk stabilizes the L1.1 region and the A site finger and the ribosomal protein uL16 the SL-III. The A site finger (28S rRNA helix 38) is a flexible component of the 28S rRNA which plays an important role in translocation of tRNAs from the A to the P site [46]. In many structures is not visible due to its intrinsic flexibility, required to perform its role escorting in-transit tRNAs [46, 47]. The SL-III of IAPV-IRES contacts the A site finger, stabilizing it in a fixed conformation, which allows the apical loop of SL-III to reach deep into the 60S, establishing a novel interaction with the ribosomal protein uL16 (Fig. 4B and C). The IAPV-IRES positions the apical loop of SL-III (nucleotides 6585 to 6590) in direct contact with basic residues of uL16, which are in electrostatic interacting distance with negatively charged phosphates of the RNA backbone of the IRES (Fig. 4C). The additional anchoring points to the ribosome contributed by SL-III, allows the IAPV-IRES, in the context of a 80S interaction, to be stabilized in a conformation that overlaps with the space occupied by a hybrid A/P-tRNA. The coaxial unit SL-III/ASL-like domain of the IAPV-IRES functionally mimmics a A/P-tRNA priming the 80S for eEF2 recruitment, effectively bypassing the initiation stage (Fig. 4D). Additionally, the anchoring points provided by the IAPV-IRES along the intersubunit space, probably contribute to an effective recruitment of the 60S in the absence of the dedicated factor responsible for such function in canonical translation, eIF5B [48].

**Figure 4:**
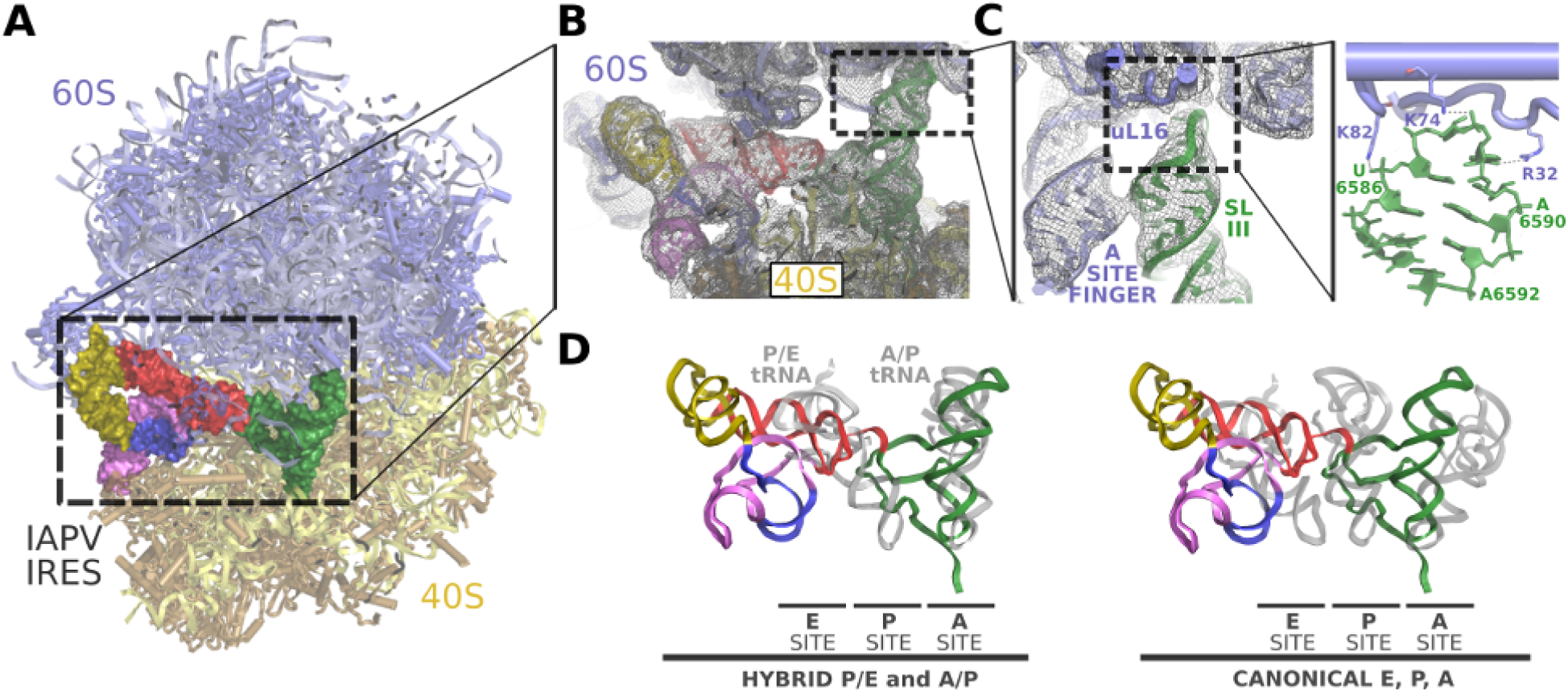
SL-III of IAPV-IRES engages novel sites of the 60S ribosomal subunit. (**A**) Overall view of the IAPV-IRES/80S complex class 1 with 60S represented as cyan cartoons, the 40S as yellow cartoons and the IAPV-IRES represented as solid Van der Waals surface colored by secondary structures motifs. (**B**) Close up view of the intersubunit space with the IAPV-IRES depicted as cartoons colored as in (**A**) inserted in the experimental, unsharpened cryoEM density. (**C**) Zoomed view of the A-site finger in interacting distance with the SL-III (green). The apical loop of SL-III reaches deep into the 60S contacting the ribosomal protein uL16. (**D**) Superposition of the IAPV-IRES in complex with 80S (class 1) with tRNAs in different configurations indicated at the bottom. Alignments were computed with the 40S body, excluding from the computation the ligands (IRES/tRNAs) and the 40S head. IAPV-IRES PKI components SL-III/ASL-like domain populate a space of the intersubunit space similar to a A/P-tRNA. IAPV-IRES PKIII (red) mimic the elbow region of a hybrid P/E-tRNA.

### Aparavirus IRESs restricts the small subunit rotation dynamics in the pre-translocation state

No populations with wide rotations of the small subunit were identified in our 80S/IAPV-IRES large dataset, what suggests that, in contrast with Cripavirus IRES’s [49, 30], the IAPV-IRES is able to restrict the dynamic of the small subunit, channeling it towards a canonical, non-rotated configuration. This is accomplished by a solid anchoring of the PKI in the A site, which not only mimics the ASL of a tRNA interacting with its cognate mRNA in the A site, but also, by placing the SL-III in a similar position as a hybrid A/P-tRNA [50, 51], mimics the T and D arms (Fig. 4D and Fig. 5B). In such position, the PKI of the IAPV-IRES establishes a network of interactions with both the large and the small subunit, effectively restricting the rotation of the 40S. Apart from the interactions stablished by the apical loop of the SL-III with ribosomal protein uL16 (Fig. 4C), the decoding event elicited by the placement of the PKI in the decoding center allows the establishment of an interaction with the 28S rRNA base A3760 (A1913 in *E.coli*), normally involved in decoding (Fig. 5C) [52]. This interaction is maintained along the small fluctuations of the 40S, which locally, are restricted to dis-placements of a few Ångströms (Fig. 5D). The IAPV-IRES seems to bind very tightly in a binary, pre-translocation complex with the 80S. The re-cruitment of elongation factors to commit the ribosome to the production of viral proteins seems to be achieved not through a dynamic manipulation of the 40S rotation, but by directly adopting a configuration reminiscent of a ribosome with tRNAs in hybrid configurations.

**Figure 5:**
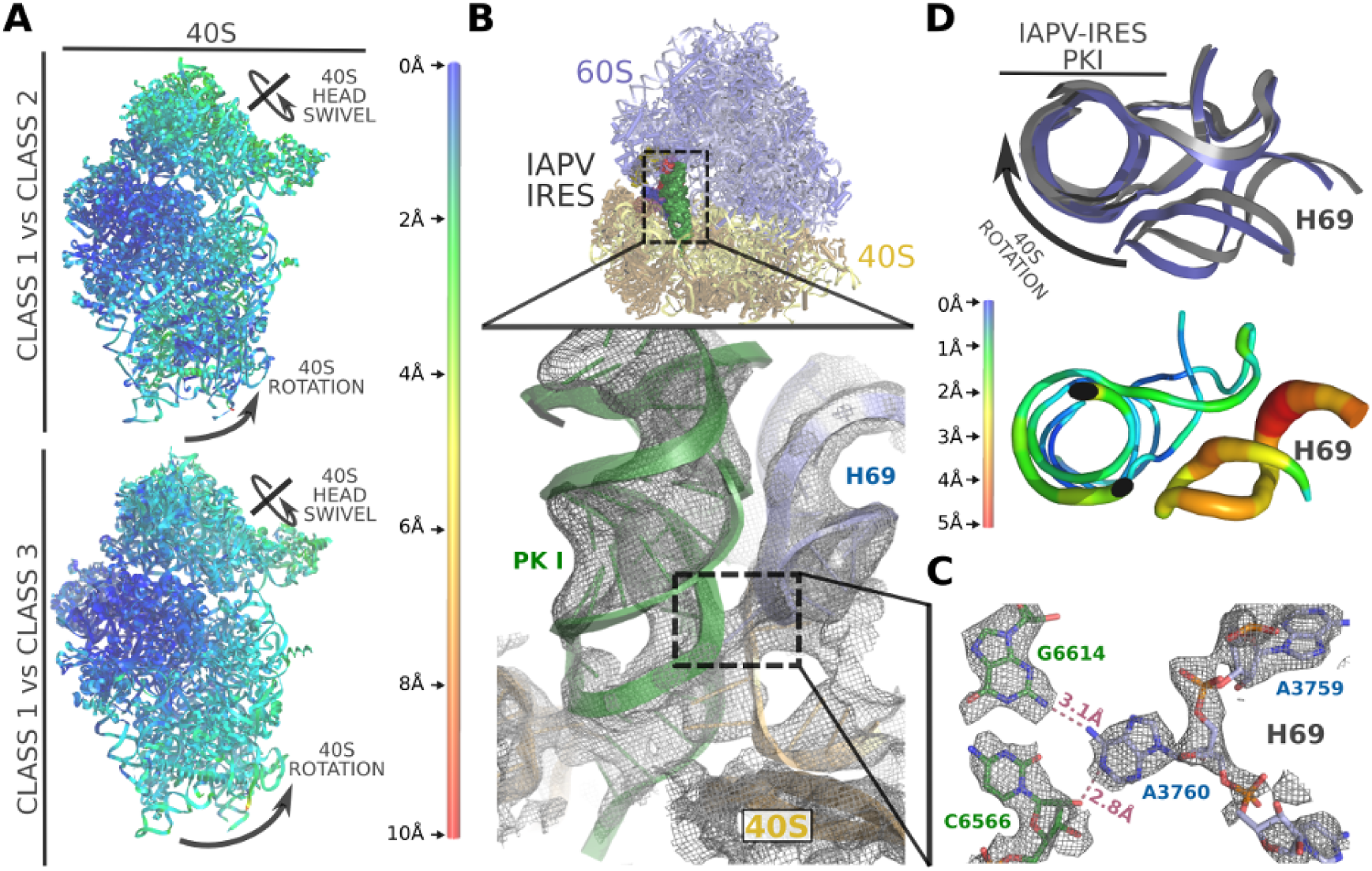
The IAPV-IRES restricts the small subunit rotational dynamic in a pre-translocation complex with 80S. (**A**) Ribbon diagram of IAPV-IRES/80S complex viewed from the 40S colored by pairwise root-mean-square deviation displacements observed between the classes indicated on the left. Class 1 is unrotated while classes 2 and 3 exhibit a small rotational movement of the 40S. (**B**) Top, general overview of the non-rotated IAPV-IRES/80S class 1 structure. IAPV-IRES is depicted as solid Van Der Walls surface colored according to the secondary structure motifs. The PKI (green) is solidly anchored to the A site. Bottom, close up view of the IAPV-IRES PKI inserted in the experimental unsharpened map. A3760, a nucleotide belonging to the helix 69 (H69) of the 28S rRNA interacts with the PKI, detail showed in (**C**). (**D**) This interaction is not disrupted along the small fluctuations of the 40S. An apical view along the axis of the PKI of a superposition of class1 versus class 2 shows the IRES displacements are minimal and are followed by the H69 which keep constantly the interaction with the IRES.

### Remodeling of specific components of the IAPV-IRES allows its translocation through the ribosome

Due to their intrinsic flexibility, IRESs are able to populate multiple conformational states, while maintaining a basic structural framework dictated by their base pairing scheme. A combination of rigid elements connected via flexible linkers allow these RNAs to tune their interactions with different ribosomal sites as they transit from an early pre-translocated state to a post-translocated one. Along this vectorial movement, these IRESs take advantage of intrinsic dynamic elements of the ribosome, normally involved in translocation of tRNAs and mRNAs [53, 23].

In order to visualize the IAPV-IRES in a post-translocated state, we engineered a stop codon in the first coding codon following the IRES sequence [26, 54]. By simultaneously incubating a pre-translocated 80S/IAPV-IRES complex with eEF2 and a catalytically inactive version of the eukaryotic release factor 1 (eRF1*) [55] in the presence of GTP, we were able to stabilize the IRES after a single translocation on the ribosome, allowing the visualization of the overall conformation of the IRES in a post-translocated state as well as the specific determinants of the IRES in binding the ribosomal P site (Fig. 6).

**Figure 6:**
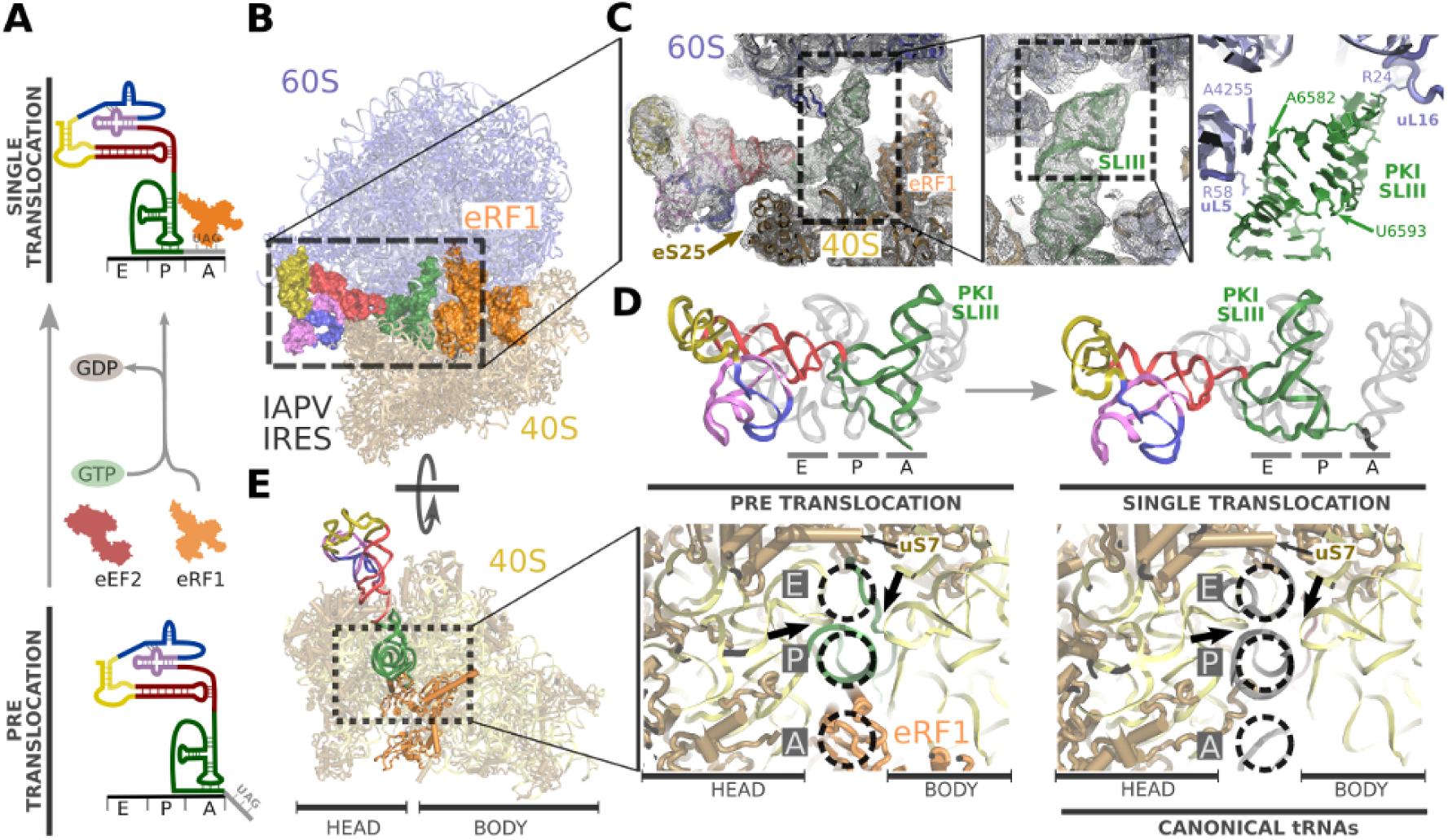
Visualization of IAPV-IRES in a post-translocated state in the ribosome. (**A**) Biochemical strategy employed to trap a single translocated state of IAPV-IRES transitioning through the ribosome. (**B**) General view of the IAPV-IRES in a post-translocated state on the ribosome: 60S depicted as blue ribbons, 40S as yellow ribbons, IAPV-IRES represented as solid Van Der Waals surface colored according to secondary structure elements described in figure 1A and eRF1^*^ depicted orange. (**C**) Close up view of the SL-III inserted in the experimental unsharpened cryoEM density. On the right, refined model with residues form the 60S (blue) in interacting distance with the SL-III (green) indicated. (**D**) Comparison of the final refined model for IAPV-IRES colored according to the secondary structure described in figure 1A with canonical tRNAs (PDBID: 4V5D) in the pre-translocated state (left) and after translocation (right). (**E**). Left, overall top view of the inter-subunit space of the 40S for the post-translocated state. Inset, close up view of the 40S tRNA binding sites where it can be appreciated the insertion of PKI of the IAPV-IRES in the P site, projecting the VRL towards the E site. The elements of the 18S rRNA forming the “P site gate” are indicated by solid arrows. On the right, equivalent view with canonical tRNAs.

Globally, the post-translocated IAPV-IRES exhibits an extended conformation with the ASL/SL-III unit deeply inserted in the P site and the L1.1 region maintaining its original connection with the L1-stalk. The SL-IV and SL-V of the IRES are no longer in contact with the 40S (Fig. 6C, left).

The SL-III seems to play an important role in orienting the IRES unit formed by the ASL/SL-III to a position that perfectly matches that of a canonical P site tRNA (Fig. 6D). Nucleotides belonging to the SL-III establish contacts with several residues of ribosomal proteins uL5 and uL16 as well as with the 28S rRNA nucleotide A4255, all components of the large ribosomal subunit (60S)(Fig. 6C, right).

In canonical translocation, the movement of tRNAs and mRNA has to be coordinated in order to vacate the ribosomal A site for the next incoming aminoacyl-tRNA. After peptidyl transfer, the ribosome adopts a rotated configuration of the small ribosomal subunit with tRNAs in hybrid configurations. The movement of the peptydil-tRNA in the A site to the P site has to be coordinated with the movement of the the P site tRNA to the E site and the tRNA occupying the E site has to be ejected from the ribosome with the assistance of the L1-stalk. During this process, is of capital importance that the correct reading frame on the mRNA be maintained [56, 23, 53]. This is accomplished by the participation of specific components of the ribosome (mainly RNA bases) which interact with tRNAs and mRNA defining specific checkpoints so as to prevent in-transit tRNAs from slipping or loosing contact with the mRNA on the correct frame [57].

One of the most important checkpoints is defined by the so called “P site gate” [57]: a constriction formed by elements of the 18S rRNA (bases 1054-1064 and 1638-1645, 1335-1344 and 785-795 in *E.coli*) that physically block the progression of a translocated peptydil-tRNA in the P site from slipping into the E site. The “closing” of the P site gate marks the end of a correct translocation cycle, allowing the ribosome to reset to a canonical, non-rotated configuration with a deacylated tRNA in the E site, a peptydil-tRNA in the P site and a vacant A site ready to accept the next aminoacyl-tRNA [56].

By perfectly mimicking a canonical P site tRNA, the IAPV-IRES in a post-translocated state, is able to position the VRL,a flexible element of its structure, in contacting distance with the E site of the 40S subunit. This is accomplish even with a fully closed P site gate (Fig. 6E): sliding through the P site gate, the single stranded VRL can contact the ribosomal protein uS7, normally involved in stabilizing E site tRNAs (Fig. 6E)[58]. The VRL thus suffers a marked remodeling as the IAPV-IRES transitions from the pre-translocation to the post-translocation state. While in the pre-translocation conformation, the VRL establishes interactions with ribosomal components of the P site, after translocation, it populates a more extended conformation able to reach the E site even with the P site gate locked. The remodeling of this IRES component seems to be crucial for proper translocation of the IRES, as mutations that alters both its length and/or its base composition impacts negatively on the ability of the IRES to initiate translation [36].

## Discussion

Metagenomic studies of environmental samples have recently underscore the pervasive role RNA viruses exerts in the biosphere [9]. With estimates of dozens of RNA viruses infecting a single species, the diversity and impact these molecular entities have in biology and evolution is highly underappreciated [59]. Insects and arthropods host the highest diversity of RNA viruses, being the *Dicistrovirideae* family of viruses of special interest given its wide range and distribution of hosts [2, 9].

Within this complex scenario, intricate population dynamics are stablished between RNA viruses, hosts and vectors, with unpredictable outcomes like decimation of entire populations, able to reshape the species compositional landscape. An example of such intricate scenario is the current collapse of bee hives worldwide, a syndrome termed <u>C</u>olony <u>C</u>ollapse <u>D</u>isorder or CCD. Even thought the exact cause of CCD could be multi-faceted and complex, there is unambiguous evidence that a related group of bee-infecting RNA viruses of the *Dicistrovirideae* family exert a deleterious effect [1, 2, 8]. Viruses of this family make use of a RNA-only mechanism to take control of host ribosomes in order to gain access to the cellular machinery for protein production [31]. Using cryo-EM, we comprehensively characterized how the intergenic IRES of IAPV, a putative agent causing CCD, binds and manipulate ribosomes to redirect them toward the production of viral proteins. The IAPV-IRES is able to establish a stable binary interaction with the small ribosomal sub-unit. An early capturing of free 40S subunits, committing them toward viral protein production, may represent a limiting step in a cellular environment where competition between cellular and viral messengers content for ribosomal access. The IAPV-IRES is able to insert its PKI domain in the A site (decoding site) of the 40S effectively blocking the binding of eIF1/eIF1A, two initiation factors required for canonical initiation (Fig. 2C). While blocking the 40S functional sites, the IAPV-IRES is able to steerer the intrinsic dynamic of the 40S subunit towards a specific configuration, facilitating the recruitment of the large ribosomal subunit (60S) in the absence of eIF5B, the cellular factor catalyzing this event (Fig. 4A).

Bypassing the highly regulated initiation stage of translation is a customary requirement for the type IV family of IRES [18]. This is accomplished by inducing an altered ribosome stage, able to recruit directly and without tRNAs, elongation factors (eEF2 and eEF1A). Cripavirus IRESs like the CrPV-IRES, accomplish this by inducing a wide rotation on the 40S, mimicking a pre-translocation state of the ribosome with tRNAs (Fig. 7, top). In marked contrast, Aparavirus IRESs like the IAPV-IRES capture elongation factors not by inducing a rotated state of the 40S, but by directly mimicking a ribosomal state with hybrid tRNAs (Fig. 4D and Fig. 7). The limited dynamic of 40S subunit rotation observed in our large cryo-EM dataset reflects a solid anchoring of both ribosomal subunits, mediated mainly by the additional contacts contributed by the SL-III characteristic of this family of IRESs.

**Figure 7:**
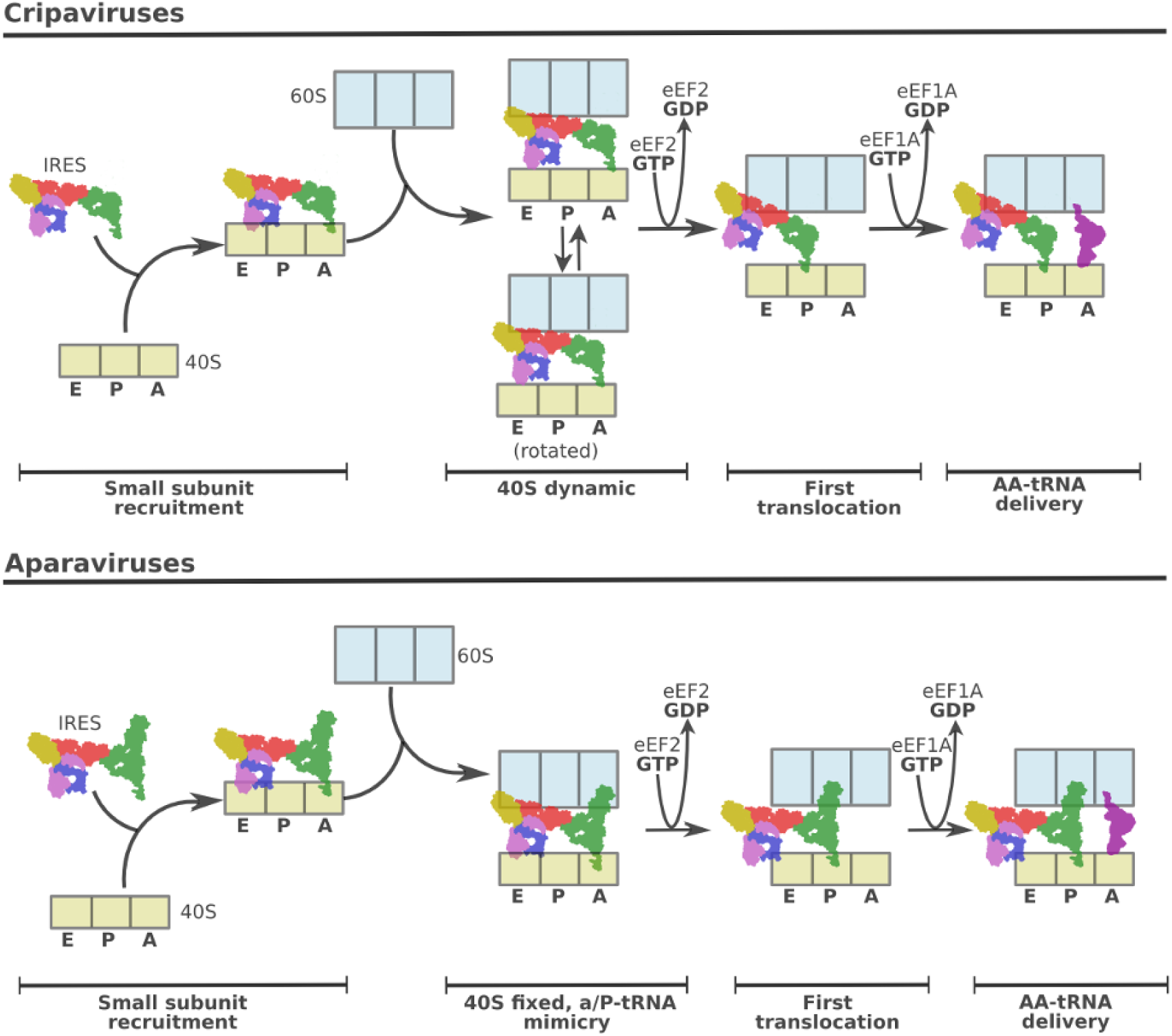
Type IV IRES families exploit different pre-translocation features of canonical translation for ribosome hijacking. (**Top**) The Cripavirus family of Type IV IRESs is able to capture free 40S subunits and engage them on a pre-translocation complex by recruiting 60S. This complex is extremely dynamic, with the 40S alternating between non-rotated and rotated configurations respect the 60S subunit. IRESs belonging to this family, exemplified by the CrPV-IRES, recruit elongation factors by mimicking a rotated stated of the ribosome with tRNAS. (**Bottom**), The Aparavirus family of IRESs follows a similar pathway in order to assemble a pre-translocation complex, however, specific structural components of this family allow for additional contacts with the 60S, limiting the rotational freedom of the 40S. Elongation factors engagement and thus effective ribosome hijacking is accomplished by mimicking a ribosome state with tRNAs in hybrid configurations.

However, a compromise between rigidity and flexibility is required for these IRESs to operate: in order to place the first coding codon in the decoding center of the 40S, the IAPV-IRES has to move (translocate) from the A to the P site. We were able to gain structural information of this transition visualizing the post-translocated state of the IAPV-IRES. In the post-translocated state, the PKI mimics a canonical P site tRNA, establishing the SL-III new contacts with proteins and rRNA components of the P site of the 60S (Fig. 6C). Additionally, featuring extremely dynamic capabilities, a single-stranded region of the IAPV-IRES termed the VRL (Fig. 2D) is able to modify its configuration in a context-specific manner. In the context of a binary interaction with the 40S subunit and in the pre-translocation state with the 80S ribosome the VRL exhibits a compact configuration, contacting bases of the 40S subunit’s P site. After translocation, and once PKI is displaced to the P site, the VRL is extended, contacting ribosomal protein uS7, a component of the 40S subunit’s E site (Fig. 6E). Biochemical evidence support a key role of the VRL in IRES function, as mutations and/or shortening impacts the ability of the IRES to efficiently translocate [36].

Using high resolution single-particle cryo-EM analysis, we showed how the prototypical Aparavirus IRES of the intergenic region of the IAPV manipulates the eukaryotic ribosome to position a viral non-AUG codon in the ribosomal A site to effectively hijack the host machinery for protein production. We have uncovered a new strategy of early pre-translocation mimicry used by this IRES sub-family as well as visualize the conformational dynamic of a strategic single-stranded segment of the IRES, the VRL. The structures presented here will allow structure-based design of new and better RNA interfering strategies directed toward the vital intergenic IRES of the IAPV. Ongoing efforts in that direction have already proved successful in fighting CCD, protecting bee hives from collapse [33, 60].

## Acknowledgements

We are thankful to Vera Pisareva and Andrey Pisarev for a generous donation of ribosomal subunits and eRF1*. We acknowledge Bob Grassucci for technical assistance in data acquisition and Harry Kao for computing. Part of this work was performed at the Simons Electron Microscopy Center and National Resource for Automated Molecular Microscopy located at the New York Structural Biology Center, supported by grants from the Simons Foundation (SF349247), NYSTAR, and the NIH National Institute of General Medical Sciences (GM103310). We are specially grateful to Ed Eng, Bill Rice and Laura Kim for support in data acquisition. Density maps have been deposited at the EMDB with accession codes XXXX and YYYY. Atomic co-ordinates have been deposited in the PDB with accession codes xxxx and yyyy. J.F. is funded by National Institutes of Health [GM097014 to J.F.] grant.

## Supplementary figure legends

**Figure S1:**
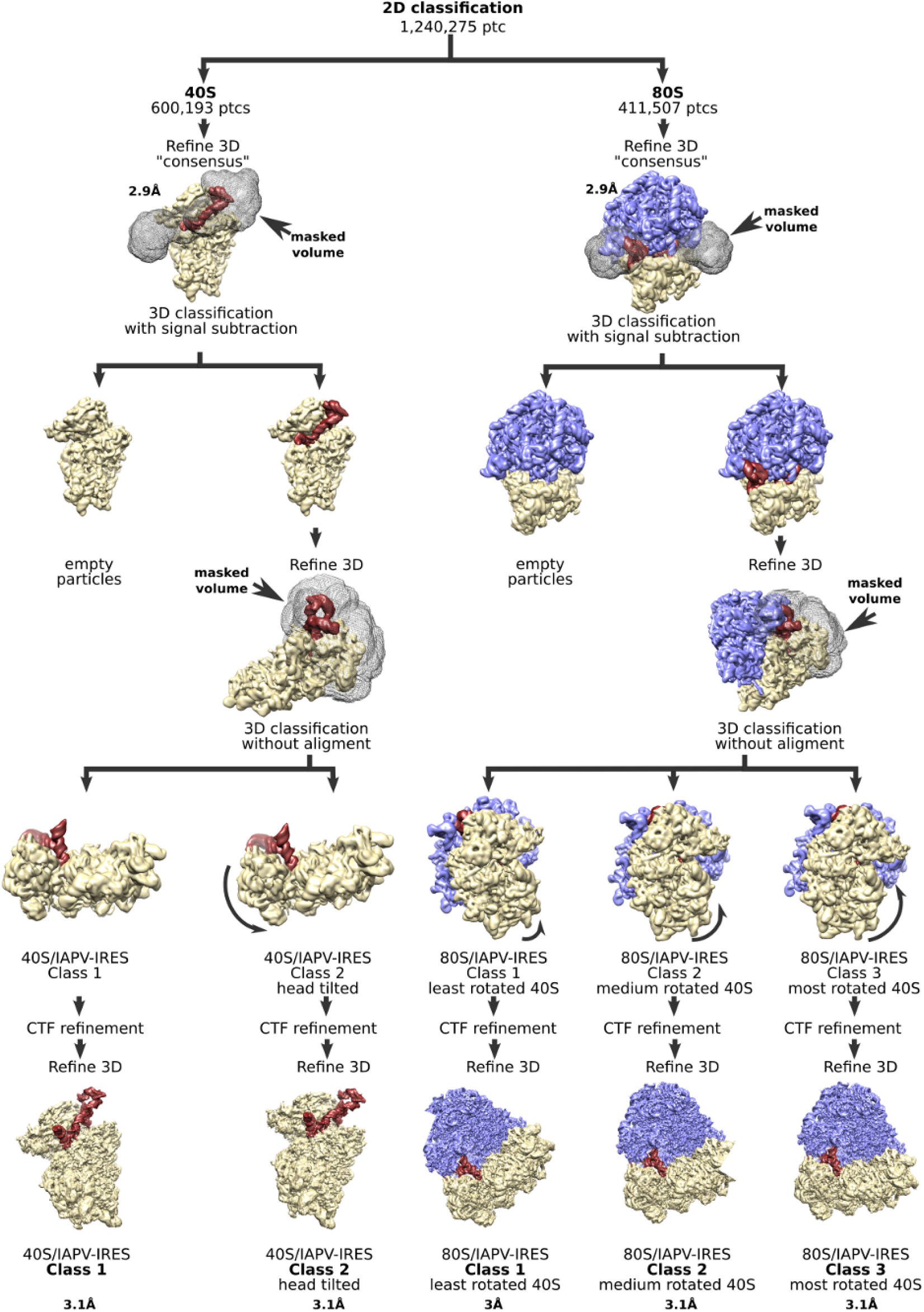
Classification scheme followed for the pre-translocation dataset. **Top**, 40S and 80S particles were separated by reference-free 2D classification alignments. Homogeneous sub-groups of 40S and 80S particles were subjected to two steps of masked classification intercalated with refinements to, on a first instance, identified those particles with IAPV-IRES and, in a second instance, distinguish among the groups with IAPV-IRES, different conformations. **Bottom** New features implemented in Relion3.0[1] such as contrast transfer values refinement allowed extending the resolution to close to 3Å for the five populations.

**Figure S2:**
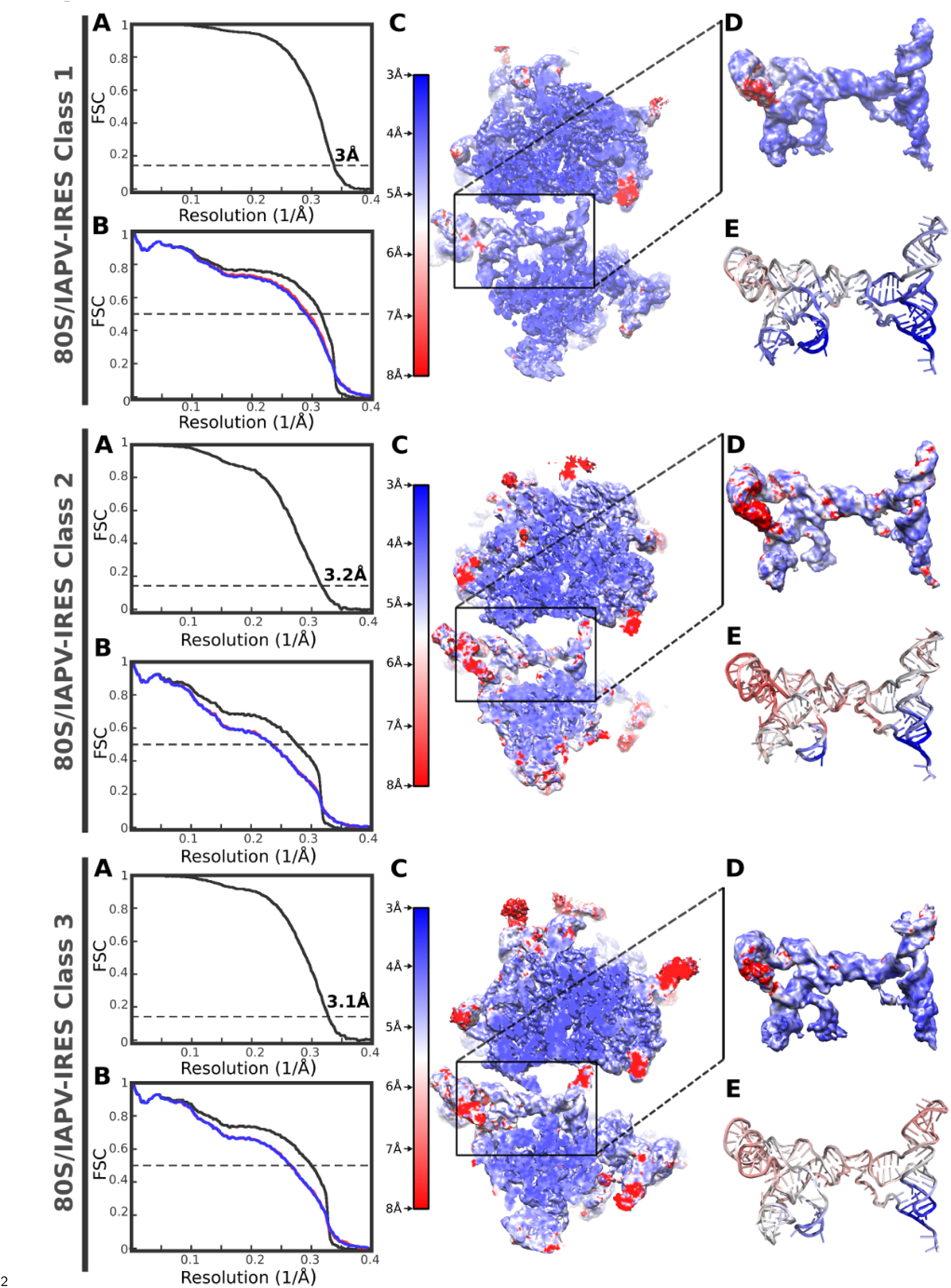
Fourier Shell Correlation curves and local resolution estimation for the 80S/IAPV-IRES complex classes. **(A)** Fourier Shell Correlation (FSC) computed for the two half maps of the final subset of particles after classification for the first class identified for the 80S/IAPV-IRES complex. The resolution is estimated to be 3-3.2Å using the 0.143 criterion[2]. **(B)** Map-versus-model cross validation FSC. The final model was validated using standard procedures: FSC of the refined model against half map 1 (blue) overlaps with the FSC against half map 2 (red, not included in the refinement). The black curve corresponds to the FSC of the final model against the final map. **(C)** Slice through the final, unsharpened map colored according to the local resolution as reported by RESMAP[3]. **(D)** Close-up views of the final density colored according to local resolution values as in **(C)** for the IAPV-IRES. **(E)** Final refined IAPV-IRES model colored according to the estimated B-factors in Å^2^ computed by REFMAC[4]. Specific values for estimated resolutions are indicated for each class.

**Figure S3:**
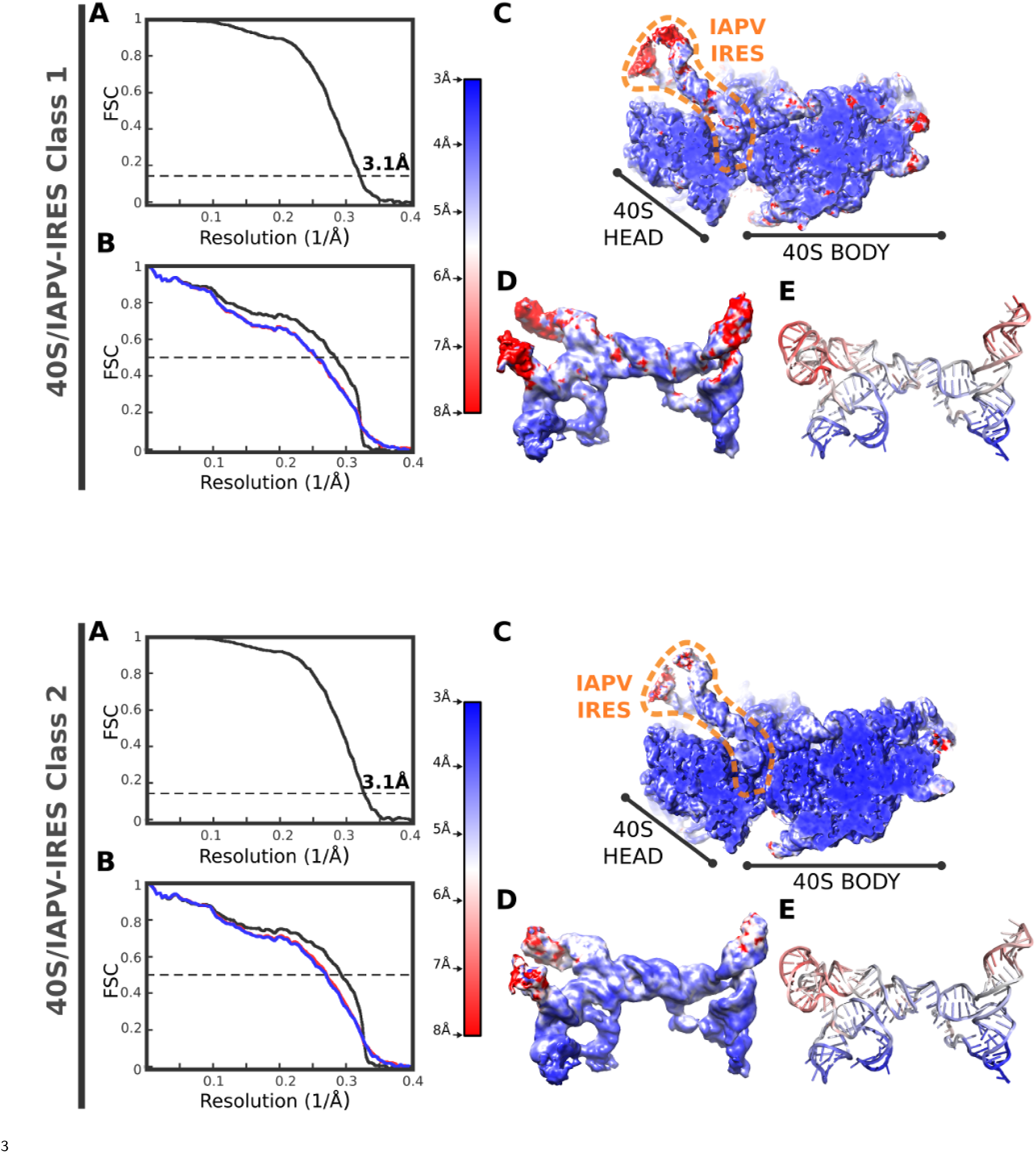
Fourier Shell Correlation curves and local resolution estimation for the 40S/IAPV-IRES complex classes. **(A)** Fourier Shell Correlation (FSC) computed for the two half maps of the final subset of particles after classification for the first class identified for the 40S/IAPV-IRES complex. The resolution is estimated to be 3.1 Å using the 0.143 criterion[2]. **(B)** Map-versus-model cross validation FSC. The final model was validated using standard procedures: FSC of the refined model against half map 1 (blue) overlaps with the FSC against half map 2 (red, not included in the refinement). The black curve corresponds to the FSC of the final model against the final map. **(C)** Slice through the final, unsharpened map colored according to the local resolution as reported by RESMAP[3]. **(D)** Close-up views of the final density colored according to local resolution values as in **(C)** for the IAPV-IRES. **(E)** Final refined IAPV-IRES model colored according to the estimated B-factors in Å^2^ computed by REFMAC. Specific values for estimated resolutions are indicated for each class.

**Figure S4:**
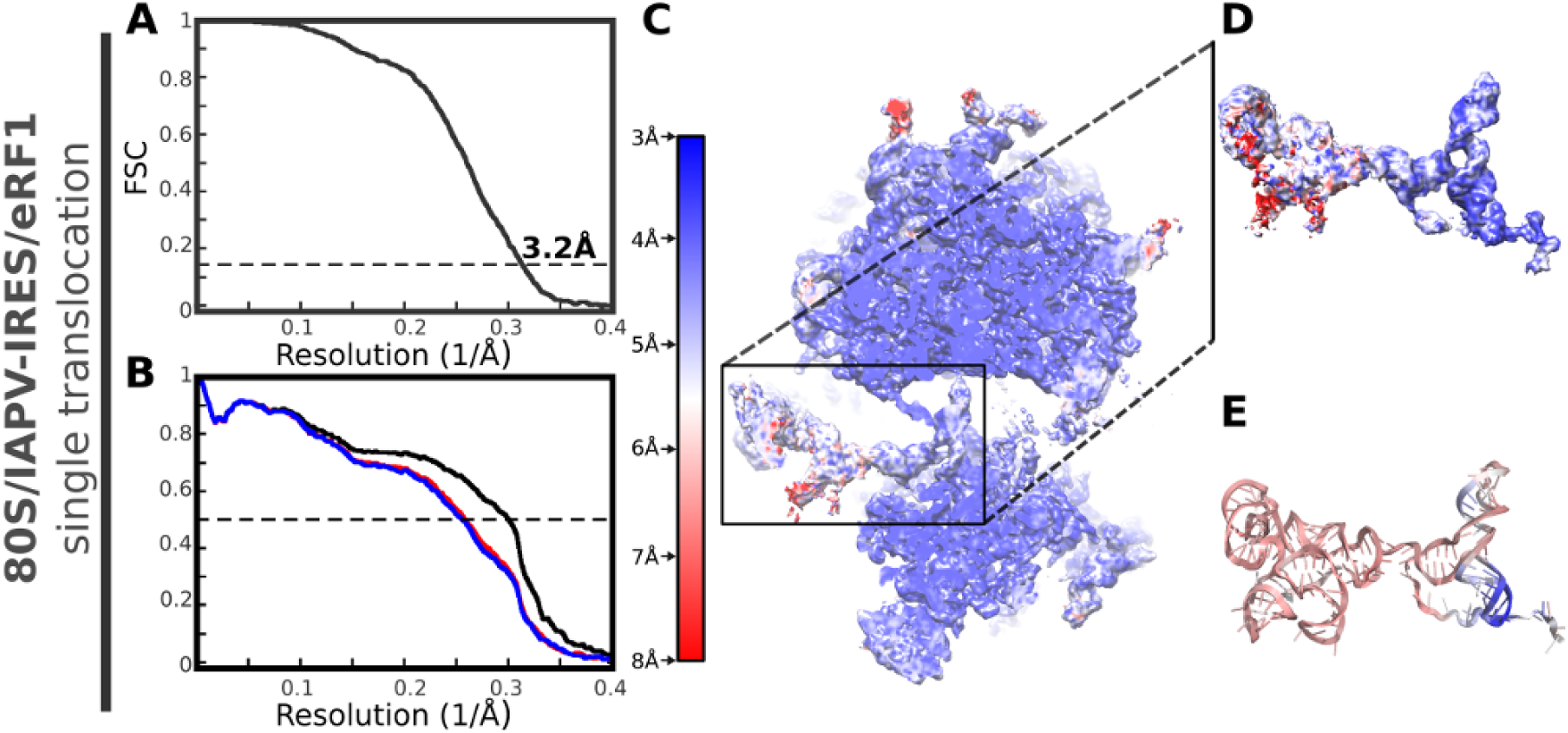
Fourier Shell Correlation curves and local resolution estimation for the post-translocated 80S/IAPV-IRES/eRF1^**\***^ complex. **(A)** Fourier Shell Correlation (FSC) computed for the two half maps of the final subset of particles after classification for the first class identified for the 80S/IAPV-IRES complex. The resolution is estimated to be 3.2Å using the 0.143 criterion[2]. **(B)** Map-versus-model cross validation FSC. The final model was validated using standard procedures: FSC of the refined model against half map 1 (blue) overlaps with the FSC against half map 2 (red, not included in the refinement). The black curve corresponds to the FSC of the final model against the final map. **(C)** Slice through the final, unsharpened map colored according to the local resolution as reported by RESMAP. **(D)** Close-up views of the final density colored according to local resolution values as in **(C)** for the IAPV-IRES. **(E)** Final refined IAPV-IRES model colored according to the estimated B-factors in Å^2^ computed by REFMAC[4].

## Movie S1: Conformational transitions of IAPV-IRES along its movement through the ribosome

IAPV-IRES binds initially to the ribosome inserting the PKI (green) in the A site mimicking a hybrid A/P-tRNA. Once translocated, PKI (green) is placed in the ribosomal P site with a configuration reminiscent of a canonical P-tRNA. Along this transition, needed to place the first coding codon of the viral messenger (black) in the A site, flexible regions of the IAPV-IRES experiment conformational transition in a context specific manner. IAPV-IRES is colored according to the secondary structure motives indicated in **Fig.1** and canonical tRNAs from PDBID 4V5D [5] are represented as semi-transparent grey cartoons. Molecular transitions have been approximated by a linear morph using Chimera[6].

**Table 1:**
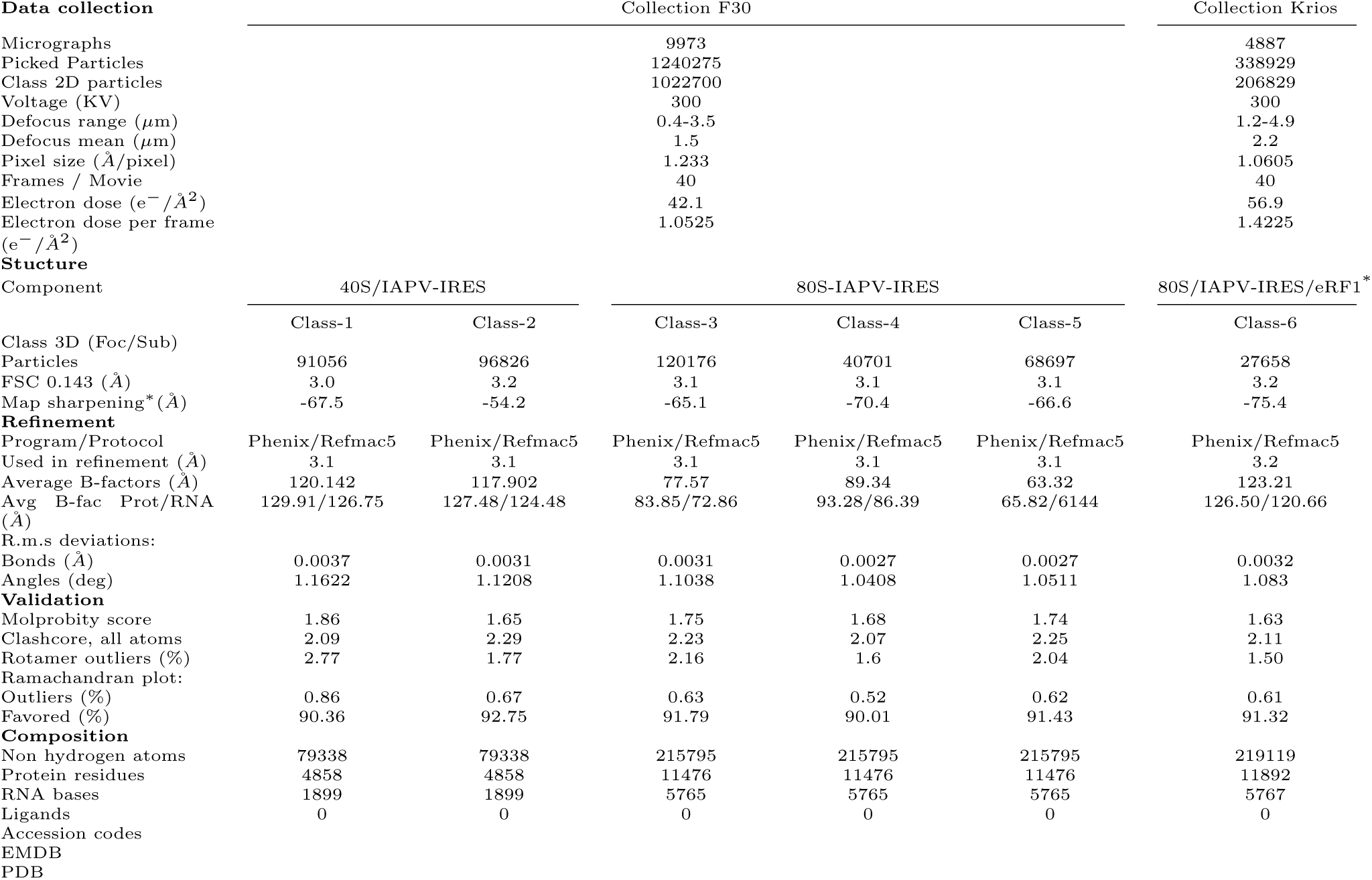
Data collection, model refinement and validation statistics.

## Materials and methods

### Plasmids

Expression vector for His-tagged eRF1*(AGQ mutant) has been previously described[1].

A pUC19 based transcription vector for IAPV-IRES-WT was constructed by inserting a T7 promoter sequence upstream of IAPV IGR IRES sequence followed by the first two coding triplets (nucleotides 6372-6623 from NC 009025). An EcoRI site was included after the second codon.

Site-directed mutagenesis was employed to change the first coding codon (GGC) to a stop codon (TAG) to create the IAPV-IRES-STOP construct. Both IRES constructs were transcribed using T7 RNA polymerase.

Briefly, 0.1 mg/mL of EcoRI linearized vector was transcribed using 0.1 mg/mL of homemade T7 RNA polymerase in 10 mL transcription buffer (100 mM HEPES-KOH pH 7.4, 10 mM of each NTP, 22 mM MgCl_2_, 50 mM DTT, 2 mM Spermidine, 1 uL/mL IPP) for 4 hours at 37 °C. The RNA was then washed, concentrated, and separated on a 6% UREA-PAGE gel. The IAPV-IRES band was cut from the gel, electro-eluted, buffer exchanged into Buffer A (20 mM Tris-HCl, pH 7.5, 100 mM KCl, 8 mM MgCl_2_, 2 mM DTT) and snap frozen in liquid nitrogen.

### Purification of translation components

Native 40S and 60S ribosomal subunits and eukaryotic elongation factor 2 (eEF2) were prepared from rabbit reticulocytes as previously described[2]. Recombinant eRF1* was purified according to a previously described protocol[3].

### Assembly of ribosomal complexes

To reconstitute ribosomal complexes in a pre-translocation state with IAPV-IRES-WT, we incubated 6 pmol of 40S ribosomal subunits with 60 pmol of IAPV-IRES-WT RNA in a 15 µl reaction mixture in Buffer A for 5 min at 37 °C. Then, the reaction mixture was supplemented with 4.5 pmol of 60S ribosomal subunits and incubated for 5 min at 37 °C. We maintained an excess of 40S ribosomal subunits to trap both the 40S-IAPV-IRES-WT and the 80S-IAPV-IRES-WT complexes in the same reaction. Assembly of the complexes was verified by running the reaction through an overnight sucrose cushion, phenol extracting the RNA and running a UREA-PAGE gel using standard methods. The assembly and integrity of the complex was also verified on a F20 screening microscope via negative stain EM and cryo-EM.

To reconstitute ribosomal complexes in a post-translocated state with IAPV-IRES-STOP, we incubated 8.8 pmol of 40S ribosomal subunits with 90 pmol of IAPV-IRES-STOP RNA in a 15 µl reaction mixture containing Buffer A with 1.7 mM GTP for 5 min at 37 °C. Then, the reaction mixture was supplemented with 8.7 pmol of 60S ribosomal subunits and incubated for 5 min at 37 °C. Next, we added 27 pmol of eEF2 and 90 pmol eRF1^*^ and incubated the reaction for an additional 30 min at 37 °C.

### CryoEM sample preparation and data acquisition

For ribosomal complexes in a pre-translocated state: 3µl aliquots of assembled ribosomal complexes at 240-390 nM concentration were incubated for 15s either on plasma-treated holey carbon grids (QUANTIFOIL R2/2 with homemade continuous carbon film estimated to be 50Åthick) or on plasma-treated holey gold grids (UltrAuFoil R1.2/1.3[4]). Grids were blotted for 2.5-3.0s and flash cooled in liquid ethane using an FEI Vitrobot. Grids were then transferred to a Polara-G2 microscope operated at 300 kV and equipped with a Gatan K2 Summit direct detector. 11,234 movies of 40 frames were collected in counting mode at 8e^-^/pix/s at a magnification of 31,000 corresponding to a calibrated pixel size of 1.25 Å. Defocus values specified ranged from 0.8 to 2.5 µm.

For ribosomal complexes in a post-translocated state: 3µl aliquots of assembled ribosomal complexes around 600 nM were incubated for 15s on plasma-treated holey gold grids (UltrAuFoil R1.2/1.3[4]). Grids were blotted for 2.5s and flash cooled in liquid ethane using an FEI Vitrobot. Grids were then transferred to a Titan Krios microscope operated at 300 kV and equipped with an energy filter (slits aperture 20eV) and a Gatan K2 Summit detector. 6904 movies of 40 frames were collected in counting mode at 8e^-^/pix/s at a magnification of 130,000 corresponding to a calibrated pixel size of 1.0605 Å. Defocus values specified ranged from 0.5 to 3.0 µm. About half the movies (3350) were collected at a 35 degree tilt to compensate for preferred orientation that was first identified during screening sessions.

On both microscopes, movies were recorded in automatic mode using the Leginon[5] software and frames were aligned using Motioncor2[6]. Data collection was monitored and checked on the fly using APPION[7].

### Image processing and structure determination

For ribosomal complexes in a pre-translocated state: Contrast transfer function parameters were estimated using GCTF[8]. For particle picking a set of templates were generated using density data obtained from a screening session on a F20 microscope that had employed Gaussian picking. Using these templates particle picking was performed using GAUTOMACH and a particle diameter value of 320 Å. The picked particles were manually screened on the micrographs to remove problematic regions. All 2D and 3D classifications and refinements were performed using RELION[9]. The picked particles were binned 4 times and subjected to a 2D classification to separate the 40S and 80S particles. We then employed 3D Refine to generate initial consensus models from both the 40S and 80S particle sets (Fig. S1). We then used our previous CrPV-IRES model (PDB: XXXX) to create appropriate masks for these initial models. The mask for the 40S initial model enclosed the putative IRES binding site. The mask for the 80S initial model enclosed the inter-subunit space, the A-site finger, the L1-stalk and ideal helices for SIII and SVI regions of the IAPV-IRES. Using these masks we performed a round of 3D classification with signal subtraction to remove 3D classes without the IRES. On the classes with the IRES, we performed focused classification without alignment using a new set of masks (Fig. S1). This focused classification step resulted in the final set of classes that were eventually used for modeling.

For ribosomal complexes in a post-translocated state we employed a similar set of protocols but with a different set of masks that also included regions for eEF2, eRF1*, and the L1-stalk in an extended conformation.

Final refinements with unbinned data for the selected classed yielded high resolution maps with density features in agreement with the reported resolution. Local resolution was computed with RESMAP[10]. New features implemented in Relion3.0[**?**] such as contrast transfer values refinement allowed extending the resolution to close to 3Å.

### Model building and refinement

Models for the mammalian ribosome and eRF1* were docked into the maps using CHIMERA[11] and COOT[12] was used to manually adjust the L1 stalk and build the IAPV-IRES using our CrPV-IRES model as initial step. An initial round of refinement was performed in Phenix using real space refinement with secondary structure restrains[13]. A final step of reciprocal-space refinement using REFMAC was performed[14] for all complexes. The fit of the model to the map density was quantified using FSCaverage and Cref.

